# Wakeful targeted memory reactivation during short rest periods modulates motor learning via the lateral orbitofrontal cortex network

**DOI:** 10.1101/2025.07.02.662893

**Authors:** Ryushin Kawasoe, Kana Matsumura, Taiga Shinohara, Koki Arima, Yuhi Takeo, Takashi Ikeda, Hisato Sugata

**Affiliations:** Graduate School of Welfare and Health Science, Oita University, Oita, Japan; Faculty of Welfare and Health Science, Oita University, Oita, Japan; Department of Rehabilitation, Oita University Hospital, 1-1, Idaigaoka, Hasama-machi, Yufu, Oita, 879-5593, Japan; Graduate School of Medicine, Oita University, 700, Dannoharu, Oita, 870-1192, Japan; Research Center for Child Mental Development, Kanazawa University, Kanazawa, Japan

## Abstract

This study investigated whether wakeful targeted memory reactivation (TMR) during short rest intervals improves motor learning. Participants were randomly assigned to the following four groups and performed a sequential key-press task under each condition: (1) *TMR_no_* group: no auditory stimuli, (2) *TMR_regular_* group: auditory cues played at the same speed as the previous task, (3) *TMR_fast_* group: auditory cues played 1.3 times faster, and (4) *TMR_random_* group: auditory cues randomized in pitch. The *TMR_regular_* group suppressed early learning gains compared with the *TMR_no_* and *TMR_fast_*groups. Electroencephalogram revealed reduced functional connectivity centered on the lateral orbitofrontal cortex (lOFC) in the *TMR_regular_* group. In contrast, the *TMR_fast_*group preserved early learning and exhibited improved lOFC-centered functional connectivity compared with the *TMR_regular_*group. Therefore, wakeful TMR might either hinder or support motor learning, depending on cue timing and structure, emphasizing the need to optimize sensory parameters for effective learning improvement.

## 1. Introduction

In our daily lives, we unconsciously engage in repetitive learning processes to improve our performance efficiency. The improvement of motor performance efficiency through repeated movement is referred to as “motor learning,” which consists of both “online learning,” during which performance improves with practice, and “offline learning,” in which improvement occurs during rest periods between practice sessions^1^. Offline learning is believed to involve memory consolidation, a process that typically entails stabilizing and improving memory traces during rest periods^2,3^. Several studies have demonstrated that memory consolidation can occur at various time points during rest periods^4–7^. Earlier studies focused on memory consolidation during sleep between practice sessions^4,5^ and demonstrated the contributions of the cerebellum, hippocampus, and striatum to memory consolidation^8,9^. More recent research reported that memory consolidation can occur not only during sleep but also during wakeful rest periods^10^. During wakeful rest, memory consolidation is induced by the primary motor cortex, dorsolateral prefrontal cortex, and orbitofrontal cortex (OFC)^7,11^.

Remarkably, studies have confirmed that consolidation can rapidly occur with only a 10-s break between motor tasks^12–14^. In the present study, skill improvement during the brief motor task is defined as “micro-online learning” and that during the brief rest period is defined as “micro-offline learning.” The experimental results revealed that “micro-offline learning” accounted for approximately 95% of the early learning gain, which has attracted increasing attention in recent years. As a neurophysiological mechanism during “micro-offline learning,” Buch et al.^13^ reported the involvement of a network architecture over several brain regions, such as the primary sensorimotor cortex, entorhinal cortex, and hippocampus. They also found that the neural activity patterns observed during the resting phase were replayed 20-fold faster than the neural activity patterns observed during the previous task performance, termed “neural replay” ^13^. In recent years, this phenomenon has been reported to be induced by the hippocampal activity in several studies^15–17^. For instance, Mylonas et al.^18^ reported that neural replay in the hippocampus is essential for procedural motor memory consolidation. Another study demonstrated that neural replay correlates with activation of the hippocampus and lateral orbitofrontal cortex (lOFC)^19^. These data emphasize that active neurophysiological mechanisms are involved during rest periods in offline learning, which play a vital role in driving improvements in motor performance.

Parallel to these data, a phenomenon known as “targeted memory reactivation (TMR)” improves learning efficiency by promoting the reactivation of specific memory traces. TMR involves associating learning content with specific sensory cues, such as auditory and olfactory stimuli, which are later presented again during sleep to facilitate memory reactivation^20–23^. For instance, Antony et al.^24^ performed an experiment in which participants listened to sounds linked to a motor task during the task, followed by the same sounds during subsequent sleep. They observed an improvement in performance accuracy in the retest compared with that in the pre-sleep test, indicating improved memory consolidation during sleep. The neural substrate of sleep TMR engages multiple brain regions^25–27^. Legendre et al.^28^ demonstrated that functional connectivity (FC) between the hippocampus and OFC is involved in TMR-induced memory reactivation, which in turn results in improved memory task performance after sleep.

In contrast to sleep TMR, several studies have focused on whether TMR improves consolidation in wakeful rest^29–32^. Veldman et al.^29^ reported that wakeful TMR accelerated the efficiency of motor learning using electrical stimuli as sensory cues. Furthermore, Salfi et al.^30^ demonstrated improvement in motor performance by TMR combined with motor imagery. These data suggest that TMR during wakeful rest improves motor performance. In contrast, Diekelmann et al.^31^ reported that a specific olfactory cue associated with a memory learning task suppresses learning by exposing it during a subsequent wakeful rest. This report suggests that applying sensory stimuli to induce TMR during wakefulness interferes with motor learning. A more recent study demonstrated that the effect of TMR during wakeful rest depends on learning contents, such as navigation and contextual learning tests^32^. Hence, there still exists a considerable debate concerning the effectiveness of wakeful TMR in facilitating motor learning. Moreover, as mentioned earlier, multiple brain regions are associated with TMR during sleep, suggesting that wakeful TMR is driven by multiple brain regions, similar to sleep TMR. Nevertheless, only a few studies have demonstrated the neurophysiological mechanisms related to wakeful TMR.

There has been extensive research in recent years on TMR incorporating sensory cues during wakefulness, and discussions are also ongoing^32^. Furthermore, studies on wakeful TMR have used auditory or olfactory stimuli ranging from several minutes to several hours^29–31^. However, the learning effects of TMR on a timescale of seconds, that is, micro-scale learning, in which rest intervals are set to a much shorter duration, have not yet been clarified in detail. Therefore, we developed the following two hypotheses: (1) applying wakeful TMR with auditory stimuli during short rest intervals further improves learning gains and (2) multiple brain regions are involved in wakeful TMR during short rest intervals.

To explore these hypotheses, we investigated how TMR affects motor learning during short 10-s rest intervals, focusing on both behavioral outcomes and neurophysiological profiles. Participants repeatedly performed a 10-s key-press task followed by a 10-s rest period. To induce wakeful TMR, task-related auditory stimuli were presented during short rest intervals, and learning efficiency was compared across four auditory stimulus conditions to examine the effects on short-term motor learning. An electroencephalogram (EEG) was recorded throughout the experiment to analyze the neural mechanisms underlying consolidation during wakeful TMR.

## 2. Results

### Changes in motor performance during short-term rest intervals

We assigned 69 participants to four groups. Five participants were excluded from subsequent analyses due to incorrect button presses during rest intervals or substantial electromyographic noise contamination in EEG data (Table 1). The experimental design consisted of practice and rest intervals for 10 s each, and participants performed a five-item explicit motor sequence task in which they pressed a key with their nondominant left hand during practice intervals, according to previous research^12^ (Fig. 1). The sequence during practice intervals was always constant (4-1-3-2-4), and participants pressed the key that matched the number as rapidly and accurately as possible. In the rest intervals, a five-item sequence was converted into “cross (×)” and continuously displayed for 10 s. In practice intervals, auditory stimuli were matched to each item with their respective musical scales and were played at the timing of the key-press. In rest intervals, the sound was played at equal intervals using the average value of the key-pressing time of the previous practice interval. Motor performance was evaluated as the correct sequence pressing speed (sequence [seq] / s).

**Table 1.**
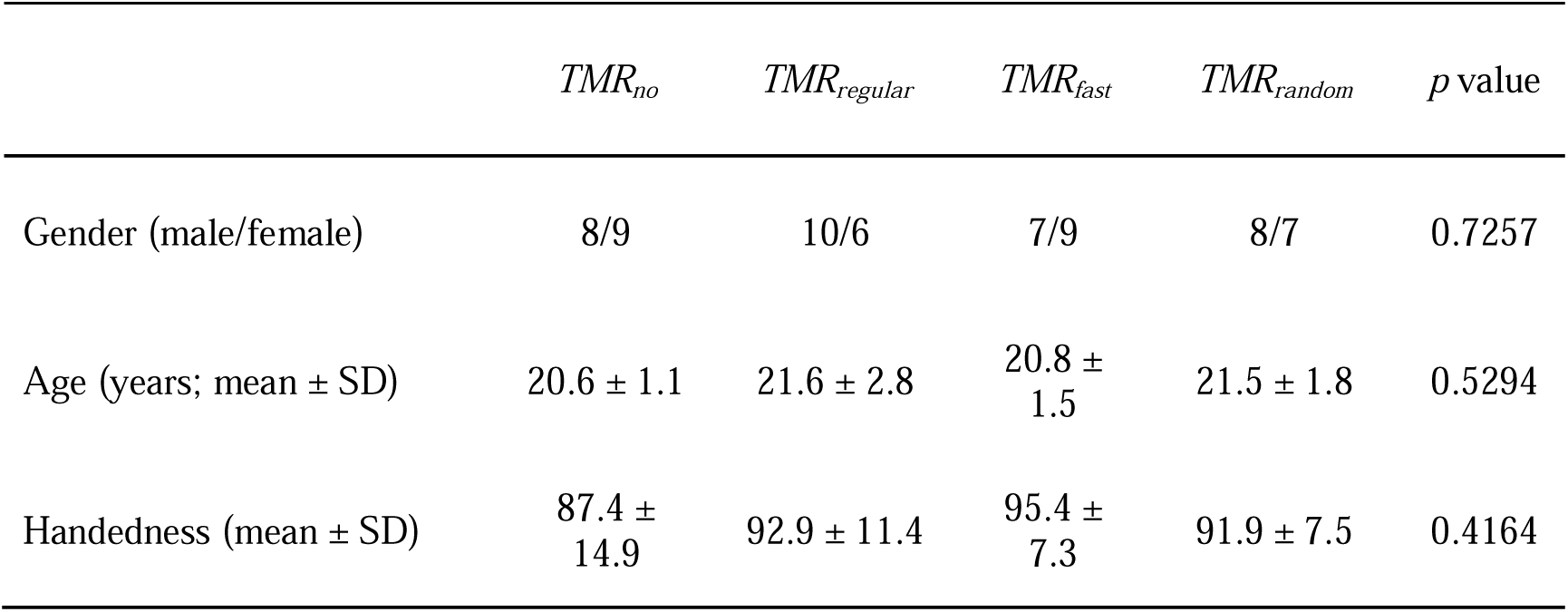
Participants’ characteristics in each group.

|  | *TMR<sub>no</sub>* | *TMR<sub>regular</sub>* | *TMR<sub>fast</sub>* | *TMR<sub>random</sub>* | *p* value |
| --- | --- | --- | --- | --- | --- |
| Gender (male/female) | 8/9 | 10/6 | 7/9 | 8/7 | 0.7257 |
| Age (years; mean ± SD) | 20.6 ± 1.1 | 21.6 ± 2.8 | 20.8 ± 1.5 | 21.5 ± 1.8 | 0.5294 |
| Handedness (mean ± SD) | 87.4 ± 14.9 | 92.9 ± 11.4 | 95.4 ± 7.3 | 91.9 ± 7.5 | 0.4164 |

**Fig. 1.**
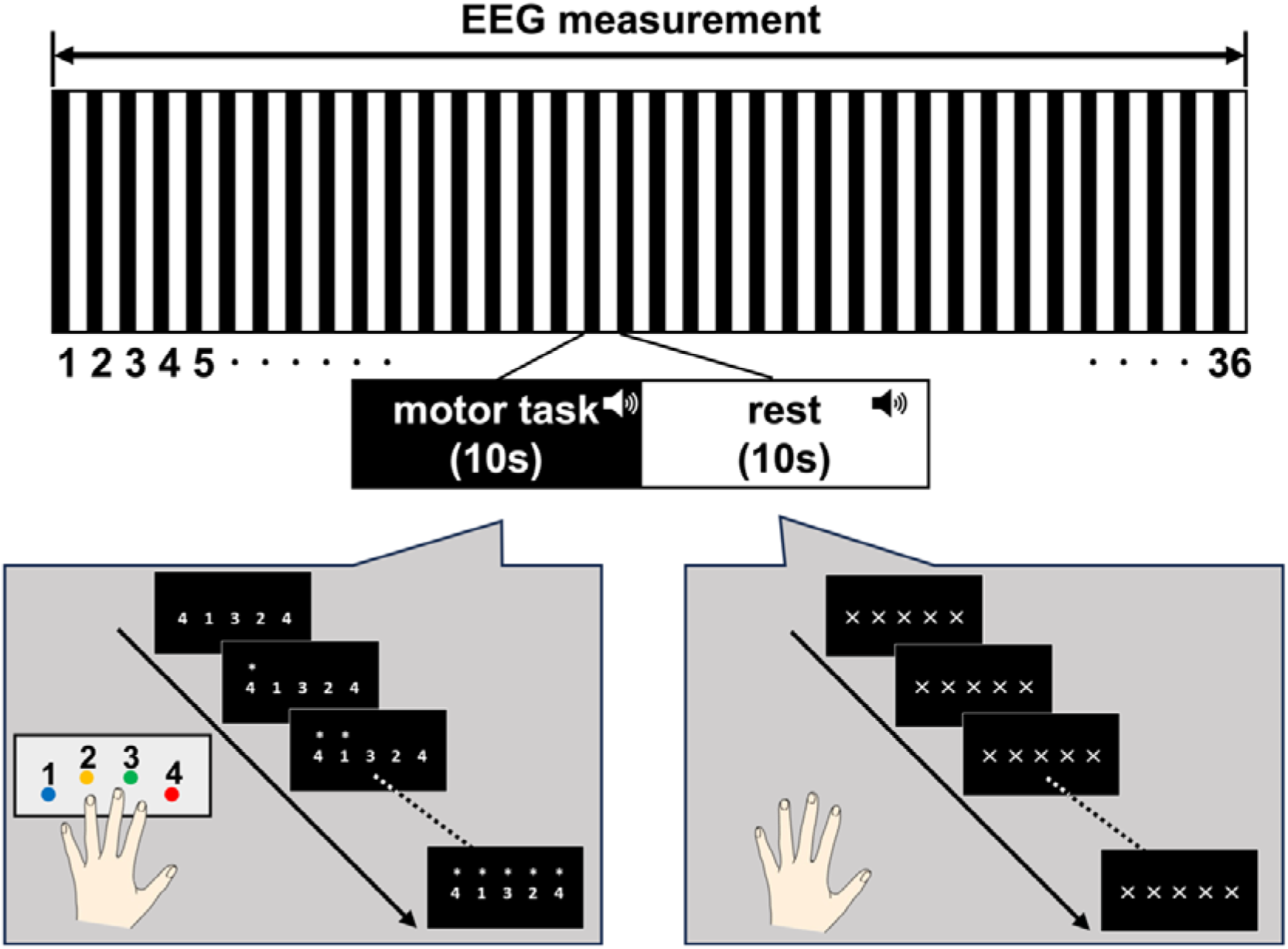
Sequential motor learning task. Each trial consisted of 10-s practice intervals followed by 10-s rest intervals, repeated for 36 trials. The motor learning task involved a key-pressing task using the nondominant left hand, in which participants were instructed to press the button corresponding to the number displayed on the screen as rapidly and accurately as possible. The five sequence items, 4-1-3-2-4, were repeated during practice intervals. The primary outcome measure was the key-pressing speed. During the rest intervals, the items were replaced with the cross (×) mark. EEG signals were simultaneously recorded throughout the experiment.

The sequence pressing speed accelerated with each successive trial (Fig. 2 A). According to a previous study, we defined learning during the practice interval as micro-online learning and learning during the rest interval as micro-offline learning^12^ (Fig. 2 B). Regarding the results, although total learning and micro-offline learning showed a significant increase in learning gain (total learning: *t* = 18.36, *p* < 0.001, *Cohen’s d* = 2.30; micro-offline learning: *t* = 6.79, *p* < 0.001, *Cohen’s d* = 0.85), micro-online learning did not (micro-online learning: *t* = −2.89, *p* < 0.005, *Cohen’s d* = −0.36, Fig. 2 C), indicating that improvement of motor performance occurred during rest intervals and not during practice intervals.

**Fig. 2.**
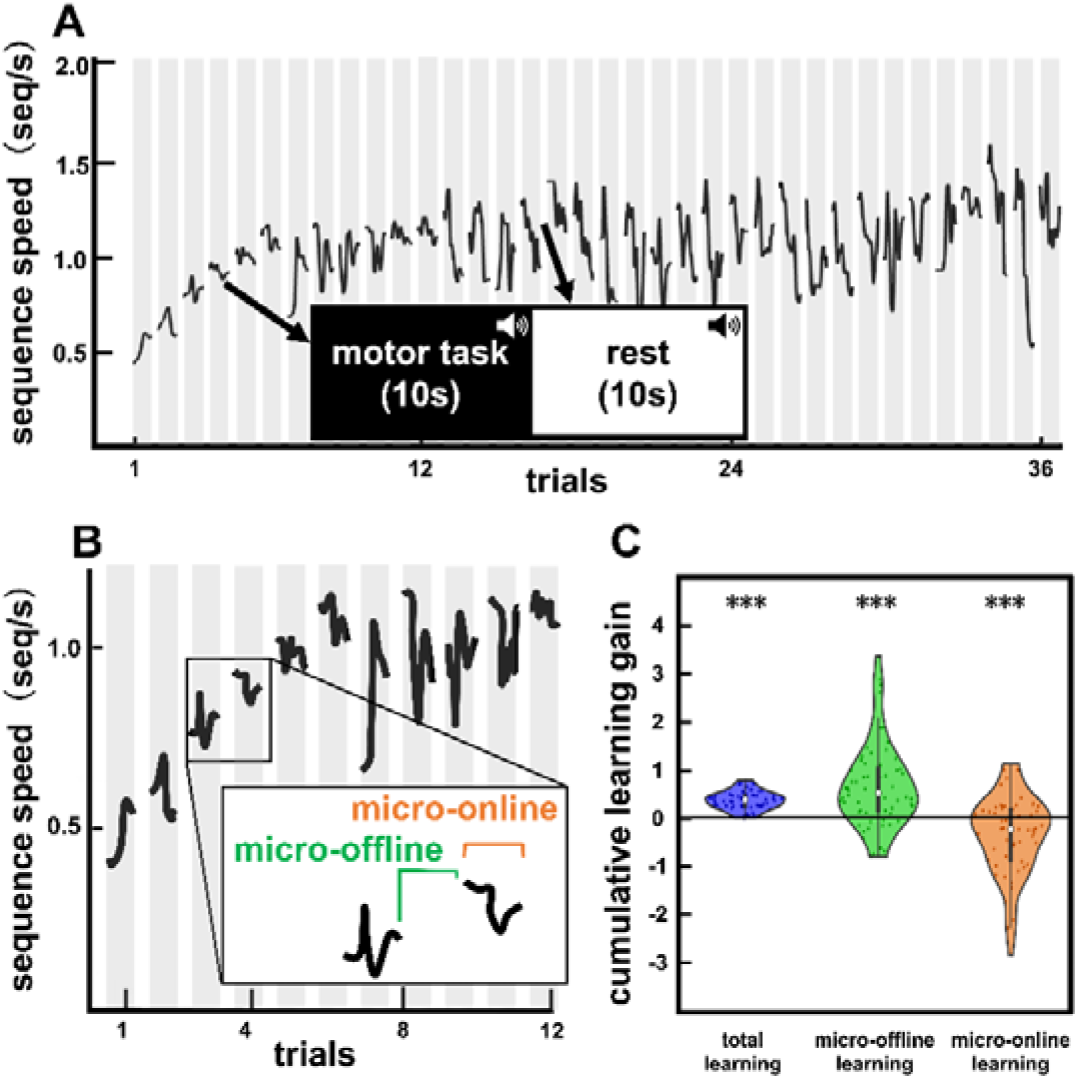
Results of motor learning task. A. Representative result of motor performance from a participant. The black line indicates the participant’s performance score during practice intervals. The vertical axis shows the sequence speed per second, and the horizontal axis indicates the number of trials. B. Schematic of a representative example in the early learning period. Micro-online indicates motor performance scores during the practice intervals, and micro-offline indicates motor performance scores during the rest intervals. C. Comparison of cumulative learning gains across all participants. The vertical axis represents the cumulative learning gain. Total learning and micro-offline learning showed significant improvement in learning gain, suggesting that micro-offline learning is responsible for most of the total learning. (***: *p <* 0.001, one-sample t-test)

### Changes in motor performance between conditions

The learning gain in each group exhibited a rapid increase in motor performance during the early learning phase (Fig. 3 A). One-way analysis of variance (ANOVA) among the four conditions revealed significant differences in early learning gain (F_(3,60)_ = 4.13, *p* < 0.01, 1]^2^ = 0.17, Fig. 3 B). The *TMR_no_* group, which did not receive auditory stimuli, showed significant increases in early learning gain compared with the *TMR_regular_* group that received auditory stimuli during the rest interval at the same speed as the practice interval just before (Fig. 3 B). Similarly, the *TMR_fast_* group, which received auditory stimuli during the rest interval at 1.3 times the speed based on a practice interval just before, demonstrated significant increases in early learning gain compared with the *TMR_regular_* group (Fig. 3 B). No significant differences were observed between the four conditions in the middle and late learning phases.

**Fig. 3.**
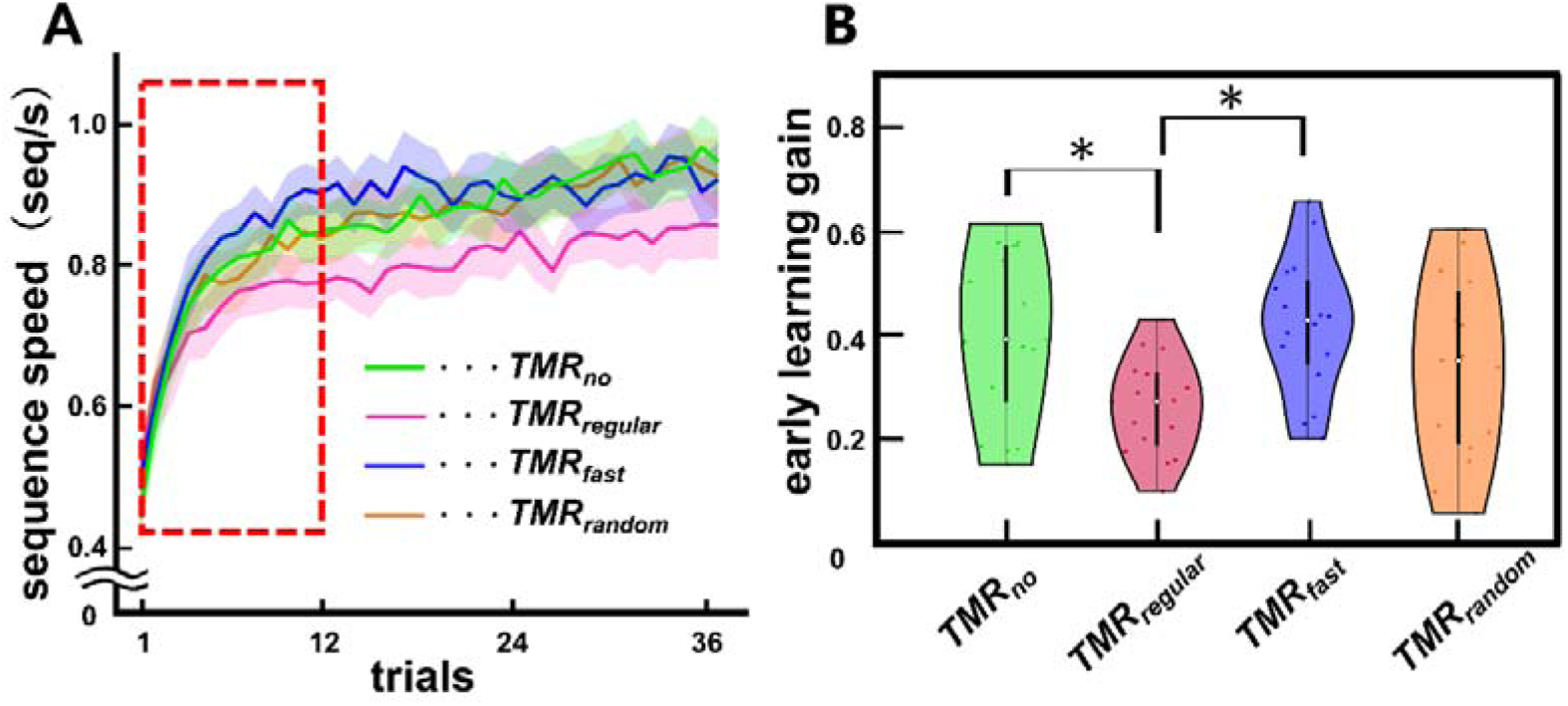
Comparison of motor performance scores. A. Improvement of motor performance in all four groups. The solid lines indicate the averaged motor performance scores across participants. Shaded areas indicate standard errors. The red dashed box indicates the early learning period during which the motor performance tends to increase rapidly. Yellow green: *TMR_no_* group, pink: *TMR_regular_* group, blue: *TMR_fast_* group, orange: *TMR_random_* group. B. Comparison of early learning gains among four groups. Early learning in the *TMR_no_*and *TMR_fast_* groups was significantly greater than that in the *TMR_regular_* group (*: *p* < 0.05, Bonferroni correction). There was no significant difference among groups in middle and late learning gain.

### Comparison of FC between conditions

To clarify the neurophysiological mechanisms related to wakeful TMR, we compared the strength of FC between groups that demonstrated significant differences in the early learning phase. In this analysis of FC, we focused on the alpha and beta frequency bands because these frequencies are related to motor learning^7,33^. We calculated FC using lagged coherence, considering the phase shifts among different brain regions^34^. The *TMR_no_* group demonstrated significantly stronger FC between the right temporal pole and left lOFC in the beta band than the *TMR_regular_* group (*p* < 0.001, *t* = 4.44, Fig. 4). Furthermore, the *TMR_fast_* group demonstrated significantly stronger FC in the alpha band between the left lOFC and right pars orbitalis (*p* < 0.001, *t* = 4.32, Fig. 5), as well as the left rostral middle frontal gyrus (RMFG) (*p* < 0.001, *t* = 4.46, Fig. 5), than the *TMR_regular_*group.

**Fig. 4.**
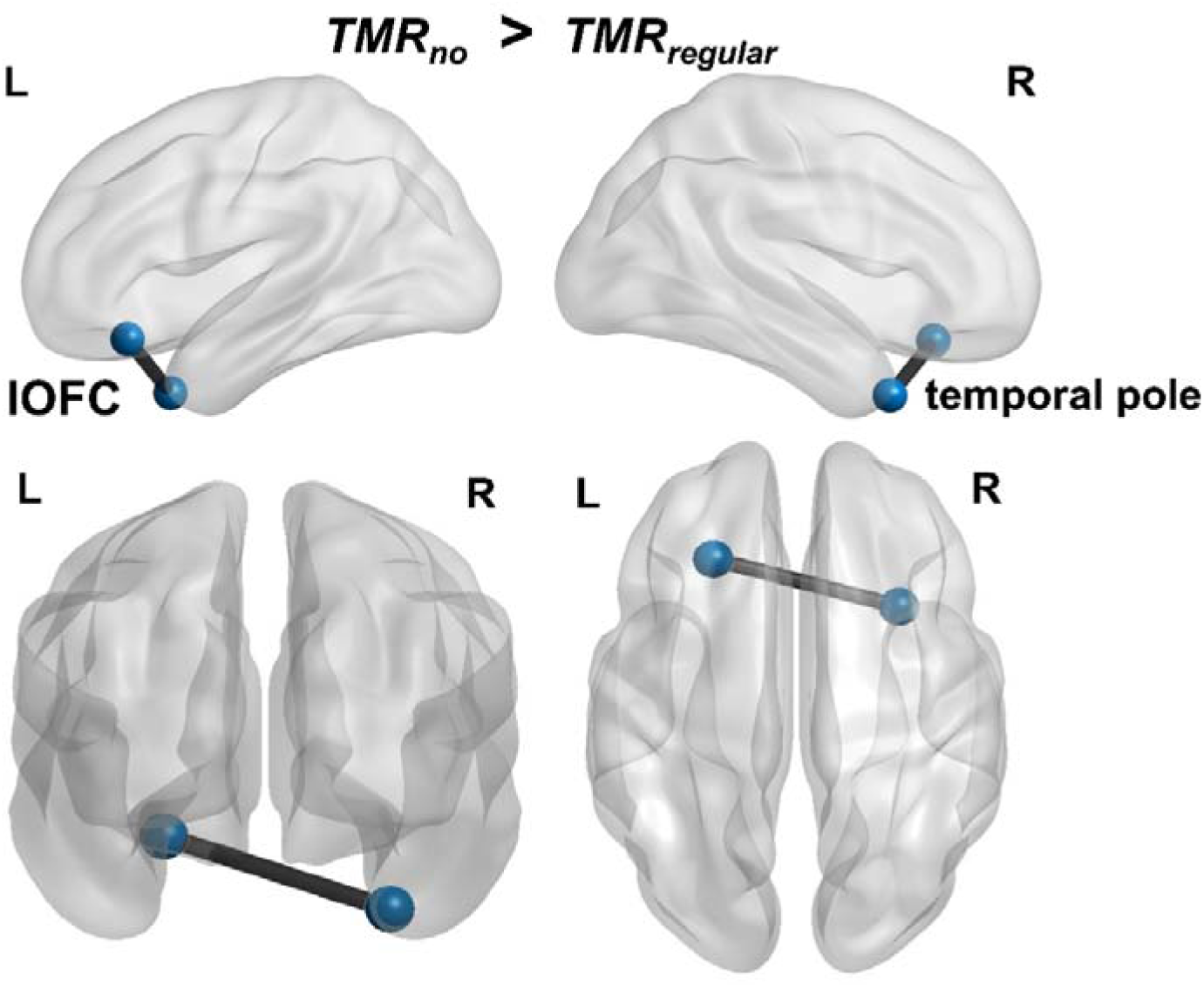
Comparison of FC between *TMR_no_* and *TMR_regular_* groups. FC between the left lOFC and right temporal pole in the *TMR_no_*group was significantly stronger than that in the *TMR_regular_* group in the beta band. L; left, R; right, lOFC; lateral orbitofrontal cortex.

**Fig. 5.**
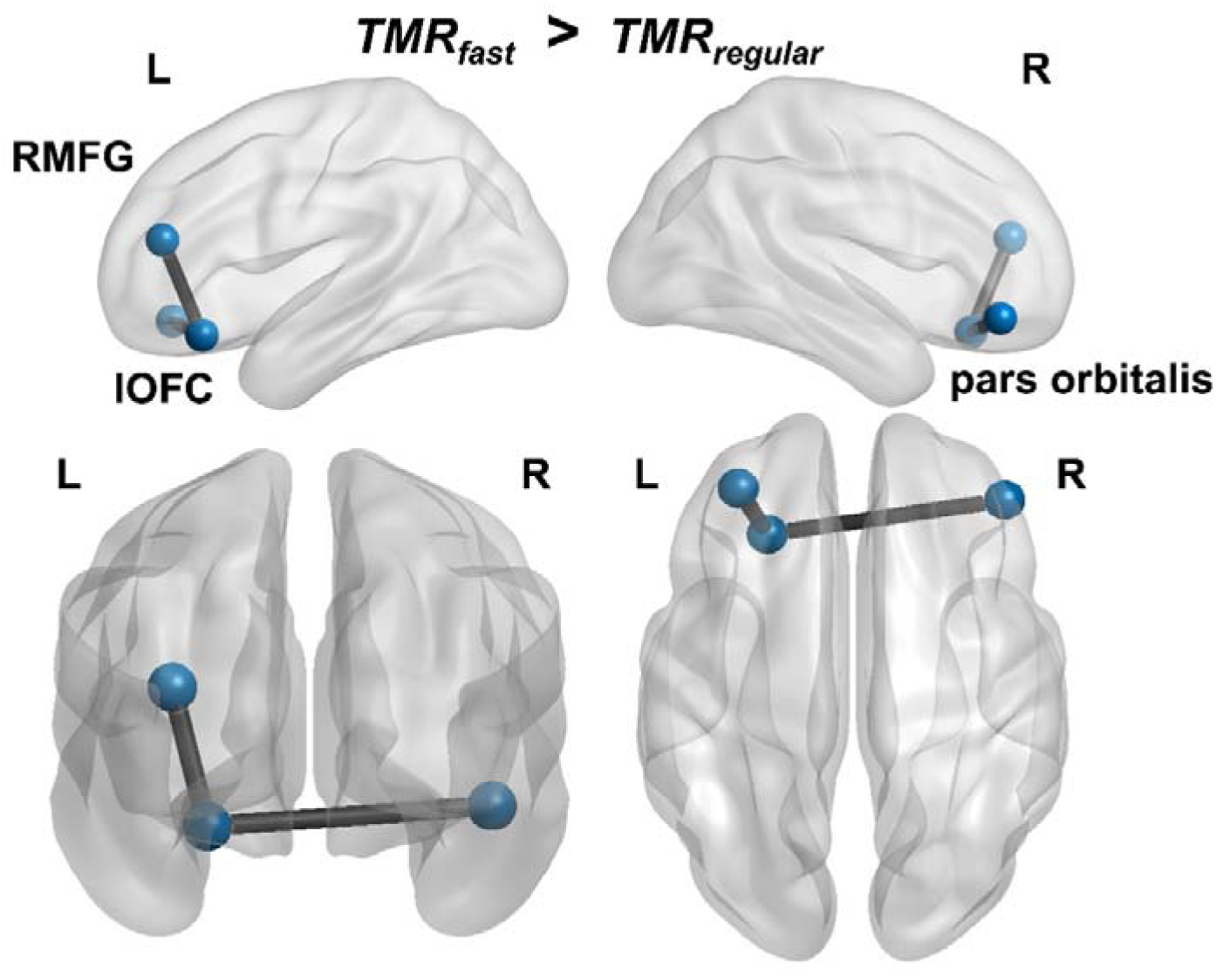
Comparison of FC between *TMR_fast_* and *TMR_regular_* groups. FC between the left lOFC, RMFG, and right pars orbitalis in the *TMR_fast_*group was significantly stronger than that in the *TMR_regular_* group in the alpha band. L; left, R; right, lOFC; lateral orbitofrontal cortex, RMFG; rostral middle frontal gyrus.

## 3. Discussion

This study explored the impact of wakeful TMR during 10-s rest intervals on micro-offline motor learning. Results showed that both *TMR_no_* and *TMR_fast_* groups had greater early learning gains than the *TMR_regular_* group, which exhibited suppressed FC centered on the lOFC.

We initially hypothesized that applying TMR using auditory stimuli during brief rest intervals improves micro-offline learning. In contrast to our expectations, the experimental results revealed that wakeful TMR inhibited motor learning.

As mentioned in the Introduction, the effect of TMR during wakeful rest on motor learning remains inconclusive. Although there is extensive evidence showing that wakeful TMR facilitated motor learning^29,30^, more recent data have suggested that wakeful TMR interferes with motor learning^31,32,35^. For instance, using task-related auditory stimuli, Hoffman et al.^35^ demonstrated that TMR during wakefulness did not facilitate early stages of memory consolidation but rather impaired it. Similarly, Diekelmann et al.^31^ reported that olfactory cues corresponding to a memory learning task improved memory function when applied in sleep conditions but declined in the wakeful rest condition. These findings suggest that TMR improves learning during sleep but inhibits it during wakefulness. A possible explanation is that, during sleep, environmental sensory input is reduced, allowing for more efficient reactivation of memory associated with sensory cues. In contrast, during wakefulness, competing external inputs may interfere with the reactivation process, thereby disrupting the early stages of memory consolidation. Altogether, our results support the notion, consistent with recent findings, that TMR during wakefulness may disturb, rather than facilitate, learning and subsequent memory consolidation.

Conversely, a comparison of early learning performance between the *TMR_fast_* and *TMR_regular_* groups, for which different speeds of auditory stimuli were applied during rest intervals, showed that the *TMR_fast_* group exhibited significantly increased early learning gain. This result suggests that the conditional settings of sensory stimuli affect early learning inhibition. In the *TMR_fast_* group, we set the auditory speed as 1.3 times the performance speed just before the rest interval, based on previous research that showed that electrical stimuli at 1.3 times the speed improved learning performance^29^. Hence, this result suggests that sensory stimuli during wakeful rest intervals do not always serve as an inhibitory factor for learning. However, it may improve motor learning, suggesting that wakeful TMR potentially results in improved early stages of memory consolidation depending on the conditional setting applying the sensory stimuli.

Supporting this notion, the *TMR_fast_* group showed no significant decrease in motor learning compared with the *TMR_no_* group, although there was no significant increase either. A possible reason for this finding is that wakeful TMR depends on the conditional setting for applying the sensory stimuli. Therefore, this result may be attributed to different task conditions (i.e., sequence type, stimulus types, and rest periods) compared with the study by Veldman et al.^29^ in which electrical stimuli were used to induce TMR. In other words, auditory stimuli at an optimal conditional setting for our key-pressing task may further improve learning performance during short-term wakeful rest intervals. Our findings suggest that applying sensory stimuli with optimal settings for inducing TMR during wakeful rest improves motor learning and subsequent memory consolidation.

Regarding the brain regions with significant FC, a comparison between the *TMR_no_* and *TMR_regular_*groups revealed that FC strength in the lOFC and temporal pole was significantly lower in the *TMR_regular_* group. These brain regions are a component of the limbic network (LIM), which is broadly constructed from the hippocampus and cingulate cortex^36–39^, along with regions such as the lOFC and temporal pole. The LIM is particularly associated with explicit motor learning^40^. Because we used an explicit key-pressing task in our study, the lOFC and temporal pole might be driven as a component of the LIM during the task. Moreover, Schuck et al.^19^ reported that the hippocampus and lOFC, which are specific components of the LIM, are deeply involved in neural replay, a phenomenon unique to offline learning. In the present study, FC related to LIM, i.e., lOFC and temporal pole, significantly decreased in the *TMR_regular_* group compared with that in the *TMR_no_*group. This result suggests that applying the auditory stimuli during rest intervals at the same speed as the practice just before had interfered with FC, including lOFC, thereby disrupting the neural replay in the subsequent offline learning process, resulting in inhibited early motor learning.

In addition, FC strengths in the *TMR_fast_* group among the lOFC, RMFG, and pars orbitalis were significantly stronger than those in the *TMR_no_* group. As mentioned earlier, the lOFC is a component of LIM; however, it has been reported that FC with RMFG is related to working memory functions involving motor actions^41^. Moreover, the FC between lOFC and pars orbitalis is related to the integration and adaptation of external feedback^42^. Several studies have also confirmed that activities in these brain regions are modulated by rhythmic auditory stimulation^43–47^. For instance, Braunlich et al.^46^ reported that FC, including the lOFC and pars orbitalis, correlated with the speed of auditory stimulation. Wang et al.^47^ also demonstrated that faster auditory stimulation results in greater activation of the prefrontal cortex, particularly the RMFG. These previous findings are consistent with our results. Our findings demonstrated that early learning was disturbed in the TMR*_regular_* group but not in the TMR*_fast_* group, suggesting that the auditory stimuli linked to task contents affect the higher order memory-related brain regions by modulating the speed of sound. These results indicate that FC among the lOFC, RMFG, and pars orbitalis was suppressed in the *TMR_regular_* group, inhibiting early learning and subsequent memory consolidation.

Our study has several limitations. First, although we used surface EEG to elucidate the neurophysiological mechanism related to wakeful TMR, it cannot reliably measure the activity of deep brain structures. Therefore, we could not examine the association of deep brain structures, such as the hippocampus, with wakeful TMR. We next applied auditory stimuli during rest intervals at 1.3 times the performance speed, based on practice intervals immediately preceding those in the *TMR_fast_* group, according to a previous study^29^. However, a previous study applied electrical stimulation and not an auditory stimulus to the fingertips to induce TMR. The electrical stimuli used in that study were directly applied to the fingertips that pressed the button; hence, the association with the learning task was direct. This difference in the modality and specificity of stimulation may explain why the modified auditory stimuli applied at 1.3 times the speed did not result in optimal improvement of motor learning and memory consolidation^12^.

To summarize, we investigated the effects of wakeful TMR on motor learning during short rest intervals. Our results demonstrated that the auditory stimuli applied at the same speed as practice intervals during the rest intervals interfered with motor learning and memory consolidation. Furthermore, this effect involves lOFC-mediated FC as a neurophysiological mechanism. Nevertheless, the inhibition of motor learning and memory consolidation may be suppressed depending on the conditional setting of the sensory stimuli. Overall, these findings suggest that optimizing sensory cue parameters during wakeful rest intervals helps mitigate learning inhibition and actively promotes motor learning and memory consolidation. Our results indicate the need to explore the optimization of stimulus conditions during wakeful rest intervals and the possibility that TMR during wakefulness can improve motor learning efficiency and memory consolidation. One such approach might be to combine TMR with other cognitive strategies. For instance, Salfi et al.^30^ reported that TMR combined with motor imagery improved motor learning, suggesting that contextual or cognitive facilitation overcomes the disruptive effects observed during wakeful TMR. This highlights a potential direction for future research aimed at refining the application of TMR in wakeful conditions.

## 4. Materials and Methods

### 4.1 Participants

A total of 69 participants were randomly assigned to four conditions for comparing the effect of TMR on auditory stimuli (mean age of participants: 21.2 ± 1.9 years; 35 men, 34 women). Three and two participants were excluded from subsequent analyses due to incorrect button presses during rest intervals and substantial electromyographic noise contamination in the EEG data, respectively (Table 1). All participants had normal visual and auditory functions and no history of neurological or psychiatric disorders. Participants were instructed to refrain from drinking alcohol the day before and from smoking 1 h before the experiment to control their physical condition. All participants were confirmed right-handed based on the Edinburgh Handedness Inventory Test^48^. Participants who had more than 5 consecutive years of piano experience in the past were excluded from the study, according to previous studies^12,13^. The experimental procedures and objectives were explained in detail to the participants, and written informed consent was obtained before the experiment. This study was conducted according to the protocol approved by the Ethics Committee of Oita University School of Medicine (Approval Number: 2606).

### 4.2 Motor learning task

We created an experimental design based on previous research to measure learning gain in short intervals using 10-s practice intervals and 10-s rest intervals as one trial for 36 trials continuously (Fig. 1)^12^. The participant sat at a desk with a monitor. In the practice intervals, the participants pressed the keypad (HHSC-1×4-CR, Current Designs, Philadelphia, PA, USA) with their nondominant left hand as rapidly and accurately as possible, corresponding to the number displayed for 10 s. The numbers displayed on the monitor consisted of five sequence items (4-1-3-2-4), and participants explicitly learned that sequence. Key-presses 1–4 corresponded to the little, ring, middle, and index fingers, respectively. An asterisk mark was displayed above the numbers at the time the key was pressed, irrespective of whether the answer was correct or incorrect. The timing (ms) of key-presses was recorded for subsequent data analysis. During the rest intervals, “×” was presented instead of the five-item sequence for 10 s. The participants were instructed to stop key-pressing and fixate their gaze on the cross mark in the center of the screen. A transition phase was set for 200 ms as a preparation period for the transition from the rest intervals to the practice intervals. The total time required for the experiment was approximately 12 min.

### 4.3 Stimulus conditions

Auditory stimuli were applied to the participants during the practice and rest intervals to analyze the effects of TMR on motor learning during short-term rest intervals. Although olfactory and auditory stimuli have often been used as sensory stimuli to induce TMR^30–32,49,50^, olfactory functions^51^ exhibit genetic variability^51^ and can be influenced by culture, environment, and daily life experienc^52^, which can cause individual differences^53^. Therefore, we adopted auditory stimuli as a sensory stimulus to induce wakeful TMR, as described by previous studies^54–56^.

In this study, we linked the key-pressing to piano sounds. Key-presses 1–4 corresponded to the musical scale “C-D-E-F,” respectively, ensuring the corresponding sound played when the button was pressed in the practice intervals. During the subsequent rest intervals, the same sounds were played to induce TMR. Even if participants pressed an incorrect item in the practice interval, corrected sounds were provided in the subsequent rest intervals, i.e., “F-C-E-D-F.” The loudness of the sound was set to approximately 70 dB, according to previous research^49^.

To demonstrate the effect of TMR on motor learning during short-term wakeful rest, we randomly assigned the participants to the following four groups:

Group 1 (*TMR_no_* group): No auditory stimulus during both practice and rest intervals Group 2 (*TMR_regular_* group): Auditory stimulus during practice and rest intervals. The sound speed provided during rest intervals matched the average key-press speed during practice intervals just before the rest interval.

Group 3 (*TMR_fast_* group): Auditory stimulus during practice and rest intervals. The sound speed provided during rest intervals was modified to 1.3 times faster than the average key-press speed during practice intervals just before the rest interval.

Group 4 (*TMR_random_* group): Auditory stimulus during practice and rest intervals. The sound speed provided during rest intervals matched the average key-press speed during practice intervals just before the rest interval; however, four types of musical scales were randomly presented.

The *TMR_no_* group was included to confirm pure micro-offline learning, and the *TMR_regular_* group was set to observe changes in learning efficiency resulting from TMR. The *TMR_fas_*_t_ group was set based on a previous study that showed that motor learning improved by applying somatosensory stimulation at 1.3 times the performance speed before learning occurs^29^. The *TMR_random_* group was included as a control group to observe the learning effects of presenting unrelated auditory stimuli.

### 4.4 EEG data acquisition

Surface EEG data were measured during the experiment to elucidate the neurophysiological mechanism of the learning effect of TMR during micro-offline learning. These data were recorded using a 64-channel EEG system (g.HIamp, g.tec, AUT) in an electrically shielded room. The EEG signals were digitally recorded using an online 0.1-to 100-Hz band-pass filter at a sampling rate of 1000 Hz. The ground electrode was located on the forehead, and the reference electrodes were mounted on the left and right earlobes. The right hemisphere uses the right earlobe as the reference electrode, and the left hemisphere uses the left earlobe as the reference electrode. The active electrodes were arranged according to the International 10-10 system. To detect eye blinks, EOG was simultaneously recorded using EEG. The electrode impedance did not exceed 50 Ω.

### 4.5 EEG data analysis

The EEG data were analyzed using the Brainstorm software^57^ implanted in MATLAB (MathWorks Inc. USA). The EEG data were subjected to independent component analysis (ICA; infomax algorithm) to detect regularly occurring artifacts such as eye blinking and cardiac response. The raw data unremovable by ICA were manually checked to remove infrequent technical and muscular artifacts and horizontal eyeball movements. Then, the EEG data were re-referenced to the arithmetic mean of all EEG electrodes (common average). Tomographic reconstruction of the data was performed by generating a three-shell sphere head model. The noise covariance matrix, including the practice and rest intervals, was calculated from the continuous EEG data. The minimum norm estimate was applied to the EEG data with 15,000 points over the cortical surface to compute the source distribution. Then, the EEG source data were parcellated into 68 regions of interest using the Desikan–Killiany Atlas^58^.

To calculate FC between brain regions, we applied lagged coherence that measures the phase congruency of two EEG signals with some phase shift^34^ because EEG between different brain regions shows a phase shift within a few milliseconds. To elucidate the neurophysiological mechanisms in micro-offline learning by TMR, we analyzed only the EEG data from the rest intervals. Moreover, we divided the 36 trials into the following three segments: early learning (1–12 trials), middle learning (13–24 trials), and late learning (25–36 trials).

Although studies have reported a relationship between motor functions and oscillatory brain activities such as theta and gamma bands^59,60^, we focused on the alpha (8–13 Hz) and beta (13–30 Hz) bands, referring to studies that demonstrated a strong relationship between motor learning and oscillatory brain activities^7,33^.

### 4.6 Statistical analysis of motor task

As mentioned earlier, we excluded three participants who pressed the button incorrectly during the rest intervals. We also excluded two participants whose EEG data were contaminated with substantial electromyographic noise during the experiment. Hence, the final sample size used for data analyses was as follows: *TMR_no_* group 17, *TMR_regular_*group 16, *TMR_fast_* group 16, and *TMR_random_* group 15 (Table 1).

Statistical analysis was conducted using MATLAB (R2022b). One-sample t-tests were performed to evaluate micro-online learning, micro-offline learning, and total learning (the combined effect of micro-online and micro-offline learning) across all groups. One-way ANOVA was performed among the four conditions to compare the effect of TMR on motor learning gains. A multiple comparison was performed using the Bonferroni correction if a significant difference was obtained among groups. Furthermore, FC strengths were compared between groups that demonstrated significant differences in the behavioral analysis using Student’s t-test.

## Data availability

Behavioral and EEG data are available upon request by contacting the corresponding author, Hisato Sugata

## Code availability

Custom-written code is available upon request by contacting the corresponding author, Hisato Sugata

## Acknowledgment

The authors want to acknowledge all volunteers who participated in this study. This study was supported by a grant from the KAKENHI [23K21590], the Japan Society for the Promotion of Science.

## Author contributions

RK and HS contributed to the experimental design, data collection and analysis, and drafting of the manuscript. TI contributed to the establishment of the motor task system. KM, TS, KA, and YT contributed to the recruitment and data collection of participants. All authors read and approved the final version of the manuscript.

